# Evolutionary strategies of *Klebsiella* phages in a host-diverse environment

**DOI:** 10.1101/2024.10.29.620800

**Authors:** Celia Ferriol-González, Pilar Domingo-Calap

## Abstract

Phage-host interactions have been studied using one or a few phages and hosts in simple, controlled systems. Here, we implemented an experimental design to evolve a 12-phage cocktail in a highly diverse environment, including a combination of 39 *Klebsiella* spp. capsular types. Our results showed how phages modulate their host range through different strategies, adopting a more generalist lifestyle or maintaining their host specificity, depending on the versatility of receptor binding proteins. Some of these proteins were mutational hotspots during the evolution experiment, mainly due to their intrinsic versatility, allowing a broader or narrower host range, and host diversity in the environment. Other mutations found in evolved phages were associated with evading bacterial defense systems or improving fitness on current hosts. Ecological dynamics between phages and hosts, including prophage activation and recombination events, also determined the fate of phage populations. In addition, single phage evolution experiments were performed to validate the results, allowing a model of receptor binding protein evolution and host range modulation to be proposed. This work is a step forward in the understanding of phage-host interactions and the appropriate implementation of phages as biomedical tools, providing insights into their impact in complex environments such as the human microbiota.

## Introduction

Viruses, as the most abundant biological entities on Earth (1), are key determinants of the ecological and evolutionary dynamics of life (2). The vast diversity and high evolutionary rates of viruses, allow their use as models to understand the mechanisms of evolution (3). Indeed, phages, viruses that infect bacteria, are particularly used for their easy handling under laboratory conditions (2,4). In nature, phages are ubiquitously distributed, and their interactions with their hosts shape the microbial communities (2,5). A major determinant in understanding phage-host interactions is the host range, defined as the diversity of hosts a phage can infect (6). Under laboratory conditions, the host range is usually assessed by testing phages in bacterial panels to measure their lytic activity, which may differ significantly from its host range in nature (2). However, it allows distinction from specialist phages with narrow host ranges (sometimes including only a single strain or a few isolates), to generalist or broad-range phages (eventually infecting different strains or bacterial species) (6). In addition, phages continually modify their host range through co-evolutionary processes with their hosts, counter-adapting to resistance mechanisms to overcome phage infection (2). Despite the low proportion of broad-range phages described using current isolation techniques, metagenomic studies suggest that host range is a continuum parameter, being broad-range phages more common than expected (2,6,7). Previous studies describe that the abundance and evolutionary advantage of broad-range phages depend on ecological parameters such as the diversity and density of the available hosts (6,8,9). Broad host range lifestyle might allow phage survival in host-diverse environments when a highly susceptible host is maintained at a low proportion. However, associated fitness costs have been described, with reduced virulence in each host. In addition, host diversity can affect adaptation given the selective pressure that maintains low fitness generalists.

Host range is primarily conditioned by the first infection step: recognition and adhesion to the receptor of the host, which is mediated by phage receptor binding proteins (RBPs) (10). Changes in the RBPs, through amino acid substitutions, or acquisition of total or partial RBPs of other phages through horizontal gene transfer, can alter host range (11,12). After, post-entry bacterial defense systems can also determine phage infection. Some examples are restriction-modification mechanisms (RM) (13), abortive infection systems (Abi) (14), regularly interspaced short palindromic repeats (CRISPRs) (15,16), and many others (17,18). Interestingly, phages have evolved diverse mechanisms to inhibit these defense systems enabling infection (19). Recognition of bacterial receptors is particularly challenging for phages infecting capsular bacteria (20). These bacteria present exopolysaccharidic capsules that protect them from diverse environmental conditions (21,22). The capsule represents the first barrier against phages, interfering in phage-host recognition by masking host receptors (20). However, some phages have evolved to overcome the capsule, including recognition of capsule receptors or degradation through depolymerases (Dpos), specific hydrolases able to recognize and digest oligosaccharide bounds of the capsule (23,24). Bacterial capsules are highly diverse (22), and the diversity of capsular types (K-types) is considered the main determinant of host tropism of phages infecting capsular bacteria (25,26). Understanding how phage tropism evolves in a highly diverse host is a challenge, and experimental evolution is an interesting tool for studying the molecular mechanisms involved in host recognition and adaptation.

Under this scenario, *Klebsiella* sp. is an excellent model of capsular bacteria. The capsule of *Klebsiella* sp. is particularly diverse, with more than 180 capsular-locus types described to date (27). Interestingly, the K-type is the main determinant of the tropism of *Klebsiella* phages, most of them being highly specific, infecting only one or two K-types (26,27). This genus is widely distributed throughout ecosystems and traditionally considered commensal, although some strains can cause human infections (28–30). Indeed, given the recent and rapid expansion of antimicrobial-resistant clones, *Klebsiella pneumoniae* has been classified as one of the most threatening pathogens worldwide (31,32). This study explores the mechanisms underlying phage adaptation in a host-diverse environment. An experimental evolution approach was used to evaluate how ecological interactions between *Klebsiella* phages and their hosts, including possible horizontal gene transfer events, influence the evolutionary process. The ancestral population consisted of a combination of phages, which evolved by modifying their host range through various strategies, reflecting both ecological and evolutionary dynamics.

## Results

### Host range increases in the presence of new available hosts

To study phage adaptation in a host-diverse environment, a 12-phage cocktail was evolved against a wide diversity of *Klebsiella* strains including 39 different K-types. The cocktail included six phages belonging to diverse families, and six with a close relative from the same genus to promote recombination (Supplementary Table 1). Regarding their host range, some phages were generalists (infecting three or more K-types), and other specialists (one or two K-types). This phage combination was previously tested against the *Klebsiella* reference strains collection (Statens Serum Institut, Copenhagen, Denmark) corresponding to the 77 reference serotypes (Supplementary Table 2), resulting in 55% of positive interactions in the 77 strains (27). In addition, an infectivity matrix of the ancestral phage cocktail containing 10^8^ PFU/mL per phage was performed against the panel of strains included in the bacterial pool used for the passages. Against the ancestral cocktail, 15/39 strains were susceptible, 18/39 strains were fully resistant, and 6/39 showed an intermediate phenotype (partially resistant), favoring phages to expand their host range due to newly available hosts during the adaptation process.

A total of 69 passages of experimental evolution were performed in three independent lineages. To evaluate phage adaptation during the experimental evolution, the host range of the evolved phage populations was tested in passages 0, 20, 40, and 69 (end of the experimental evolution). The number of susceptible strains increased over time, achieving the broadest spectrum in passage 40 of lineage 1. During the experiment, 10 strains with an ancestral phenotype resistant or partially resistant became susceptible. To validate how phages adapted to the experimental conditions the host range was also tested against the entire reference collection, which includes the 77 serotypes. Interestingly, a reduction of infectivity was observed in the strains not included in the evolution, validating that phages tend to specialize in preferentially infecting available hosts. The most dramatic reduction occurred in passage 69 of lineage 2, where 7 strains with a susceptible or partially resistant phenotype became fully resistant (Fig. 1, Supplementary Table 3).

**Figure 1.**
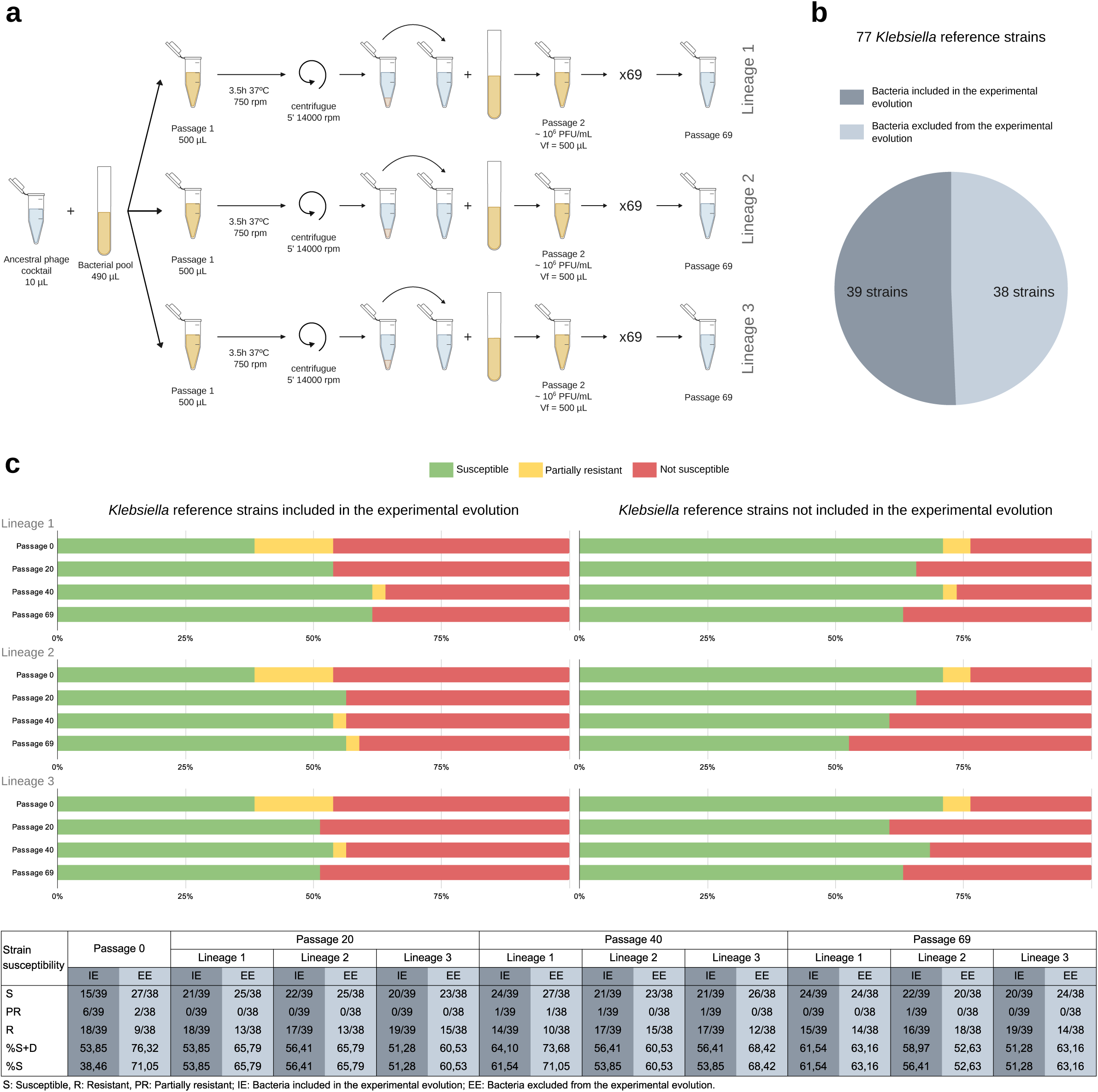
Experimental setup and host range evolution of the viral populations. **a.** Experimental setup. The ancestral 12-phage cocktail (10⁸ PFU/mL per phage) was combined with a bacterial pool and evolved during 69 passages in three parallel lineages. **b.** Composition of the bacterial pool from the 77 *Klebsiella* reference strain collection. 39 strains were included in the pool corresponding to 39 different K-types. The whole collection was used to evaluate variations in the host range of evolved phage populations. **c.** Evaluation of host range of the different lineages. Samples were tested at passages 0, 20, 40, and 69 of the three parallel lineages. The graph on the left represents the percentage of K-types included in the bacterial pool that are susceptible, resistant, or partially resistant. The graph on the right shows these data for the reference K-types not included in the bacterial pool during the evolution. The table summarises proportions and percentages of K-type strains susceptible, resistant, and/or partially resistant.

### The presence of alternative hosts enhances the parallel accumulation of mutations in RBPs

After host range evaluation, the evolved phage populations were sequenced at passages 40 and 69 (ancestral phages previously sequenced (27)). Moreover, single plaques observed in new hosts (strains not infected by the ancestral cocktail) were isolated by plaque-to-plaque purification and sequenced (Supplementary Table 4). Genomic data revealed that, despite being large double-stranded (ds) DNA phages, they accumulated mutations over time, some being adaptive. Variability accumulated mainly in RBPs, showing differential patterns between specialist and generalist phages.

The specialist phages K44PH129C1, K60PH164C1, K5lambda5, K74PH129C2, and K8PH128, encoded between 2 and 5 RBPs each (27). For K44PH129C1 (*Vectrevirus*), K60PH164C1 (*Gamaleyavirus*), and K5lambda5 (*Taipeivirus*), the only susceptible strain included in the bacterial population was their isolation strain (K44, K60, and K5 respectively). Mutations detected in these phages rarely surpassed allele frequencies higher than 0.5 and were consistent through lineages, and any variant was isolated in new hosts. For K8PH128 and K74PH129C2, both *Przondoviruses* encoding 3 RBPs (27), their isolation strains, and other partially susceptible hosts were included in the bacterial pool. In the analysis of K8PH128, parallel mutations in the RBP2 and in a serine-threonine kinase were detected (Fig. 2A, Supplementary Table 5). Again, any variant of this phage was isolated from newly susceptible strains. In the case of K74PH129C2, 6 variants were newly isolated in K80. The analysis showed 14 mutations in 10 CDSs, three of them non-synonymous in the RBP2 (Supplementary Table 6). Interestingly, RBP2 of both phages was a tail spike protein with a right-handed β-helix Dpo domain (Fig. 3) (27). The N-terminal domain of RBP2 of both *Przondovirus* was 90.07% identical (E-value: 1e-104) in 98% of query coverage (Supplementary Table 7) and the rest of the protein sequence diverged, suggesting previous RBP domain swapping by horizontal gene transfer between these phages in nature (12).

**Figure 2.**
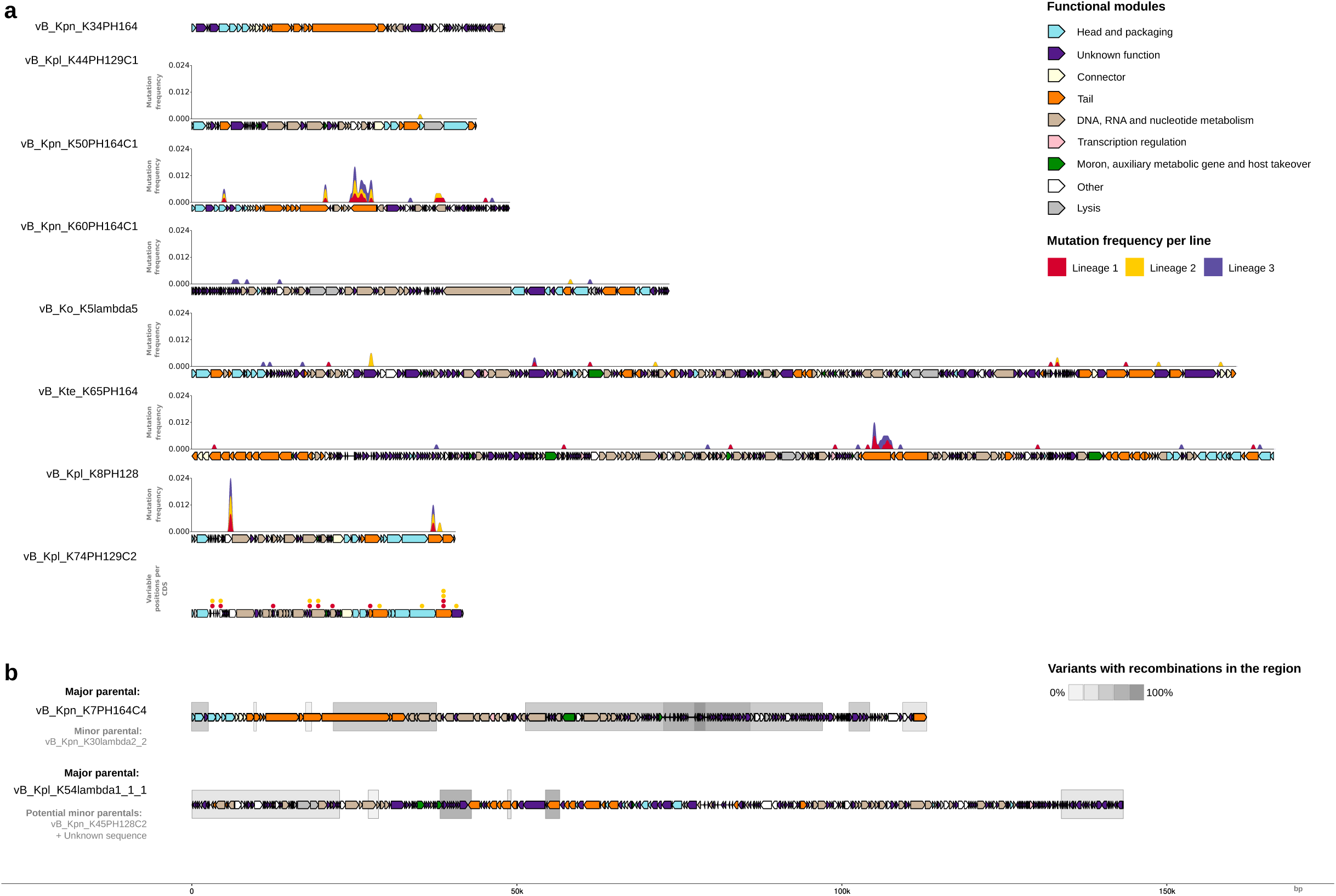
Genomic annotation and mutations accumulated in the evolution experiment for each phage. **a.** K34PH164 was lost in the three lineages. For phages K44PH129C1, K50PH164C1, K60PH164C1, K5lambda5, K65PH164, and K8PH128 the graph represents the proportion of mutations using sliding windows of 500 bp obtained from variant calling data. For phage K74PH129C2 variable positions per CDS detected in the whole genome alignment of isolated variants of this phage are represented with dots. Each dot represents a point mutation in the CDS. Colors of curves and spots correspond to the lineage where mutations were detected. **b.** Phages affected by recombination events. Only the major parent is represented. Gray squares represent the relative number of isolated recombinants that present a recombination event affecting the region. Representation of the genomes was done using R package gggenomes (70).

**Figure 3.**
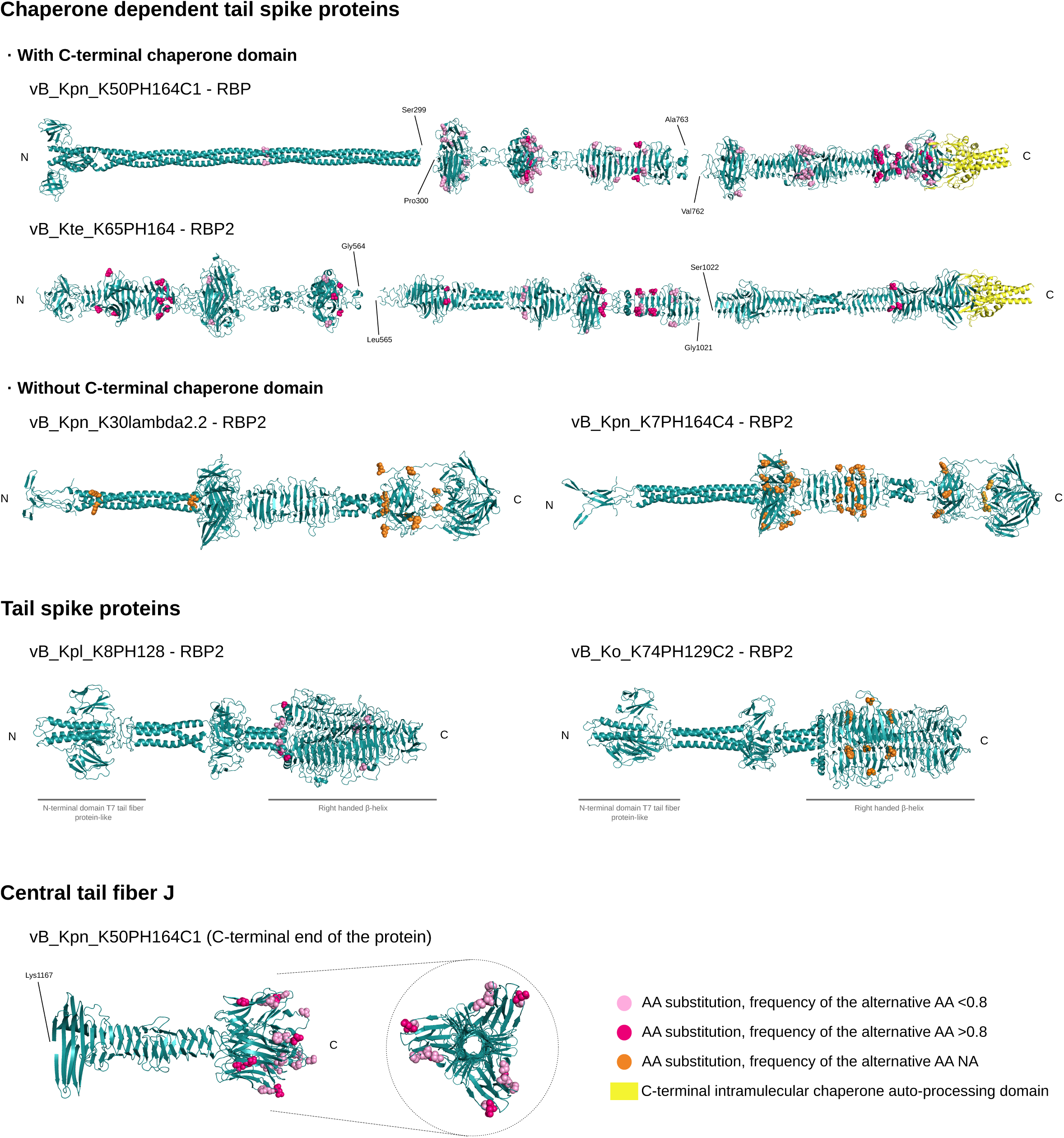
**Folding and variable positions of the proteins involved in host recognition and attachment affected by parallel evolution**. The 3D structure was obtained using AlphaFold Server (AlphaFold3 (79)) from the protein sequence of the ancestral phages. Amino acid changes detected in the protein alignment of the multiple variants are represented as spheres. For those phages where the variant calling analysis was available, those positions where the alternative amino acid was at higher frequencies than 0.8, the spheres were colored in fuchsia, while when it was lower, they were pink. Variable amino acid positions in phages where the variant calling analysis was unavailable are indicated as orange spheres. Variant positions of protein RDP2 of K8PH128 were those detected using the variant calling analysis due to the lack of isolated variants. The C-terminal chaperone auto-processing domains are colored in yellow. C: C-terminal end, N: N-terminal end. AA: amino acid. NA: Non-available.

Phages K65PH164 (*Jiaodavirus*) and K50PH164C1 (unclassified *Drexlerviridae*) were considered generalist phages. Both encoded a chaperone-dependent tail spike with a C-terminal intramolecular chaperone auto-processing (ICA) domain and a triple-helix Dpo domain that showed a parallel accumulation of single nucleotide substitutions (SNPs) at high frequencies (RBP2 of K65PH164 and RBP of K50PH164C1) (27) (Fig. 2A, Supplementary Table 2 and 7). Throughout the passages, 10 and 38 evolved variants of K65PH164 and K50PH164C1 respectively were isolated infecting novel hosts (Supplementary Table 4). Whole genome alignment corroborated a common hotspot in the RBPs (Supplementary Table 6). Amino acid changes were majorly distributed through the β-sheets and loops of the Dpo of both RBPs, without affecting the chaperone cap. In addition, for phage K50PH164C1, the central tail fiber protein J (27) showed variable positions in the C-terminal end (Fig. 3, Supplementary Table 8). Other mutations in individual evolved variants were also observed (Supplementary Table 6). The isolation and characterization of single plaques allowed us to differentiate patterns of amino acid changes in RBPs with an impact in the host range. To evaluate this, 4 and 6 evolved variants of phages K65PH164 and K50PH164C1 respectively were selected. Their host range was assessed, showing an increase in the host range for some variants, while a decrease or slight variation for others. For K65PH164, the variant with the broadest host range was K65_evo1. When evaluating those phages on strains not included in the evolution, K65_evo1 improved its infection ability in ancestral hosts. The other variants showed a more specific phenotype, losing their infection ability in some strains. Regarding variants of K50PH164C1, the only newly infected strain was K62 by K50_evo36, which loses its infection ability in K53, a highly susceptible host for the ancestral phage (Supplementary Table 9). A total of 6 variants were isolated in K62, but no common mutation between them was found (Supplementary Table 8).

To evaluate if the accumulation of RBP variability was dependent on the abundance of alternative hosts, we quantified the number of different variable positions per lineage detected in the variant calling analysis (VPL) (Supplementary Table 5). A higher number of VPLs means a higher number of polymorphic positions. We observed higher variability in RBPs of generalist phages, which may also target more K-types in the pool than the RBPs of specialist phages. To confirm our results, a second evolution experiment was performed. Five phages were evolved in solitary against the bacterial pool during 20 passages: K50PH164C1 (generalist), K65PH164 (generalist), K8PH128 (specialist with two hosts in the pool), K5lambda5 (specialist with one host in the pool), and K60PH164C1 (specialist with one host in the pool). Three parallel evolution lineages were carried out independently for each phage and every final passage was sequenced. Variant calling analyses revealed a similar variability pattern compared with the evolution of the cocktail, validating our hypothesis (Supplementary Table 10).

### Phages with close relatives in the cocktail tend to recombine

The cocktail design included genomically closely related phage pairs to enhance recombination. Phages K7PH164C4 and K30lambda2.2 are broad-range *Sugarlandviruses* with an intergenomic similarity of 90.3. During the passages, 27 recombinants were isolated, being the major parental K7PH164C4 for all of them. Most recombination events overlapped in a region including CDSs with unknown functions (Fig. 2B, Extended Data Fig. 1). Interestingly, all included the CDS 98 of K30lambda2.2, which is a hypothetical protein with a GIY-YIG family endonuclease domain, related with invasion behaviors (33).

Additionally, some recombination events affected recognition proteins. Both phages encoded RBPɑ (putative tail Dpo with 6-bladed β-propeller Dpo domain), RBPβ (tail fiber protein), and RBPγ (chaperone dependent tail spike with triple-helix Dpo domain); as well as a central tail fiber J (27,34). RBPγ was the most divergent between the parental phages (78.92% of identity). For the 27 recombinants, this RBP was almost identical to K7PH164C2 in 20 of them (100-99.71% identity, 100% query, E-value: 0.0) and to K30lambda2.2 in the other 7 (100-99.41% identity, 100% query, E-value: 0.0). Curiously, all phages isolated in K47 were recombinants with the RBPγK30lambda2.2 (Supplementary Table 4), suggesting its role targeting this K-type. Additional single mutations in the RBPγ were widespread along the β-helix region of the protein (Fig. 3). Regarding the other recognition proteins, changes observed were not associated with the selective infection of any specific K-type (Supplementary Table 8).

### Phages can optimize their life cycle by deleting large fragments of their genome

The isolation of plaques from passages allowed a detailed characterization of the evolution of the single genomes. Interestingly, 6 isolated variants from phage K74PH129C2 reported large deletions affecting similar regions. A 749 bp deletion was present in all the isolated variants (deletion 1, positions 3826-4576 bp in the ancestral phage). In addition, a 647 bp deletion was detected in all variants isolated from lineage 1 and 2/4 from lineage 2 (deletion 2, positions 9637-10285 bp), and an 1174 bp deletion was found in the other 2/4 lineage 2 variants (deletion 3, positions 9620-10794 bp, including deletion 2). CDSs affected by deletions were a SAM-dependent methyltransferase (deletion 1), a dGTPase inhibitor (deletion 3), and others with unknown functions. These proteins are usually involved in overcoming bacterial anti-phage defense systems: methyltransferases allow the escape from RM systems (35), while dGTPase inhibitors may prevent replication inhibition by bacterial dGTPases (36,37). The loss of these genes during the experiment suggested they were unnecessary for phage K74PH129C2 to replicate in its available hosts (Supplementary Table 6).

To validate if deletions affected phage fitness, we compared the infectivity of the ancestral K74PH129C2 and two evolved variants of this phage with different patterns of deletions. Phages were adjusted to an initial titer of ∼ 10^7^ PFU/mL to infect K74 at a MOI (multiplicity of infection) ∼ 0.1. Phage growth was significantly higher for evolved phages than for the ancestral one (ANOVA, Tukey’s multiple comparisons of means, p-value: 4.8e-07) (Extended Data Fig. 2), demonstrating that large deletions increased phage fitness in the conditions tested.

### Prophages activate during the experimental evolution

*Klebsiella* spp. usually encode prophages in their genomes, which can be activated under experimental conditions (38,39). Thus, the *Klebsiella* strains used in this study were screened for prophage fingerprints. Genomic analyses of the passages revealed the activity of multiple prophages in strains K15, K38, K61, and K74, without a common pattern of activation between lineages (Supplementary Table 11). In addition, other viral contigs that did not share significant homology with our bacterial genomes were detected, probably belonging to bacteria with unavailable genomes, as they had the highest similitude with viral contigs obtained in metagenomic analyses (40). Finally, some contigs corresponding to putative prophages were identified in passage 0, belonging to strains used to amplify ancestral phages (previously to cocktail elaboration) (Supplementary Table 11).

We further evaluated the prophage activation in the experimental evolution of individual phages. First, the host range revealed a very similar pattern among all, independently of the phage evolved. Interestingly, contigs with >99% identity (E-value: 0.0) to a K20 plasmid were found in all lineages (Supplementary Table 12). This plasmid is similar to other *Klebsiella* plasmids, but also to *Klebsiella* plasmid prophages vB_Kpn_1825-KPC53 (99.19% of identity, 90% query coverage, E-value = 0.0) and ST13-OXA48phi12.3 (99.18% of identity, 71% query coverage, E-value = 0.0) (41). To validate whether this was a plasmid prophage responsible for the common host range pattern, we analyzed bacterial cultures in the absence of phages. The supernatants of 4 hour post amplification cultures were assessed for all the conditions, confirming the common host range pattern (Supplementary Table 13).

### Lytic phages can recombine with putative prophages from bacteria

K54lambda1.1.1 and K45PH128C2 are *Mydoviruses* with an intergenomic similarity of 83.7 encoding 6 and 9 RBPs respectively (27) and were expected to recombine during the experiment. However, only 6 isolated evolved phages were similar to these *Mydoviruses*. Interestingly, recombination events were detected, revealing K54lambda1.1.1 as the major parental for all recombinants, with several recombinant regions of unknown origin, suggesting recombination with a putative prophage (Supplementary Table 14, Extended Data Fig. 3). A region affected in all six recombinants was located between positions 54341-55297 bp (beginning breakpoint range in all variants) and 56767-58301 bp (ending breakpoint range), encoding for a tail spike protein with a right-handed β-helix Dpo domain. This RBPK23 was 99.96% identical in 100% of the query to a tail spike protein of vB_Kpn_K23PH08C2 (E-value: 0.0), and >97.4% similar in >99% of the query to a putative tail spike protein of phages KpS8 and vB_KpnM_Seu62 (E-value: 0.0). All these are *Mydoviruses* isolated in *Klebsiella* strains with K-type 23, suggesting this tail spike is involved in K23 infection (27,42). Variability in other RBPs of the major parent K54lambda1.1.1 was also analyzed. Mutations and recombination events detected in these proteins are detailed in Supplementary Tables 8 and 14, but none of them were associated with the selective infection of a particular K-type.

### Ecological interactions between phages and bacteria determine phage survival

The analysis of evolved populations provided a better understanding of the ecological dynamics that determine the survival and abundance of phages. The genomic data revealed the loss of some phages during the evolution. Two phages were lost in only one lineage: K74PH129C2 in lineage 3, and phage K65PH164 in lineage 2. The absence of this event in other lineages may indicate that their loss was stochastic. In addition, phage K34PH164 was lost in all three lineages. Their available hosts were K60, K68, and K69, all of which were susceptible to other phages present in the cocktail (K60PH164C1, K30lambda2.2, and K7PH164C4 respectively), which could compete with K34PH164. The ubiquitous loss of K34PH164 suggests that it plays a disadvantageous role in the competition for hosts. For K60, K34PH164 competes with a specific phage better adapted to this host. For K68 and K69, however, it competes with generalist phages. These phages have many other alternative hosts in the pool, being able to acquire a larger proportion of the population more quickly, which may favor them in the competition and displace K34PH164, the less efficient competitor. Thus, K34PH164 may be disadvantaged in the competition for all its available hosts, which leads to its loss in all lineages.

Ecological dynamics were also observed among the bacterial population thanks to the independent phage evolution experiments. As previously reported, differences in the prevalence of bacterially expressed prophages are conditioned by the availability of hosts where the prophage can complete its lytic life cycle, which, in turn, is determined by the presence of different lytic phages in the system and host-host interactions. The profile of active prophages detected in cocktail evolution differs from the single-phage evolution experiments, the most obvious difference being the detection of the K20 plasmid prophage. The strains included in the bacterial pool in which the K20 prophage can replicate were K44, K60, K62, K67, and K69. All of these strains were susceptible to the initial phage cocktail. For this reason, host availability for K20 prophage replication during cocktail evolution would be lower than during single phage evolutions.

The correlation between the abundance of a particular phage and its hosts can be illustrated by the dynamics of phage K5lambda5. This phage was detected in all cocktail-evolving lineages at titers close to 10^6^-10^7^ PFU/mL, and at least one order below in single-phage evolving lineages. In fact, in the single-phage evolution experiment, K5lambda5 was not detected by sequencing. A possible explanation could be the slower growth of phages in the system due to low host abundance in the pool or interactions with other bacteria, which may interfere with each other while growing. These results support the interest of ecological relationships in modulating populations and phage-host relationships.

## Discussion

The host range in phages is a continuous parameter that varies from broad to strictly narrow, influenced by ecological factors such as the diversity and density of hosts (6). An interesting model to study how this continuum evolves is *Klebsiella* since it has been demonstrated that commensal and pathogenic strains can coexist within patients (43). The intricate nature of *Klebsiella* populations heightens the interest in studying their interactions with phages, especially in diverse and fluctuating environments. Here, a combination of *Klebsiella* phages was experimentally evolved in a host-diverse environment to provide further insights into phage evolution and ecological dynamics. Our findings revealed that phage populations tend to adapt to available hosts, but ecological relationships between hosts and phages create a complex framework for understanding their adaptive mechanisms. A single phage can take on either a specialist or generalist role based on the availability of hosts and the presence of competitors, as previously reported (44). Moreover, bacteria can compete between them, which may favor the growth of certain bacterial strains over others (45), impacting the phage population. At the same time, prophages can activate and infect other bacteria, affecting host availability and promoting recombination (46). These factors and others (including stochasticity) can significantly influence phage evolution.

Previous work of a single phage evolved in a host-diverse environment showed the selection of generalists versus specialists (8). Moreover, it has been described that in a context that favors host range expansion, host recognition proteins can accumulate mutations appearing to be mutational hotspots (47). Indeed, a few amino acid changes in the RBP can modify the host range (8,48,49). Our experimental setup allowed us to go further by including phage diversity as a variable, as occurs in nature. First, we observed the adaptability of *Klebsiella* phages through modifications at the RBPs, as previously (26,27). However, other factors are affecting phage evolution. Based on our observations, we propose a model for the molecular evolution of RBPs in host-diverse communities including two main scenarios: in the absence and the presence of recombination (Fig. 4). We consider the potential or cumulative host range of a given phage as the sum of the host ranges of the evolved variants and their ancestor. All the hosts a phage can infect with a few amino acid substitutions are considered potentially susceptible hosts. In addition, there is a gradient in the versatility of the RBPs. For versatile RBPs, the cumulative host range is broad and can differ for each variant’s host range. In contrast, specific RBPs confer a narrow host range that rarely differs from their cumulative host range, showing less versatility. The intrinsic versatility of an RBP and the availability of different potentially susceptible hosts in the environment determine the degree of variability that the RBP will accumulate. Indeed, in narrow-host range phages, depending on whether there is a single high-infectivity host or an alternative low-infectivity host available, variability in RBPs is absent or limited. In addition, our experimental conditions allowed co-infection of different phages, thanks to the overlapping host range of some of them. In this case, a phage can acquire RBPs through recombination with other lytic phages, but also with prophages. Acquired RBPs can be more or less versatile, and the accumulation of different RBPs with complementary specificity has been proposed as a fast strategy for host range expansion (50,51). Our proposed model of RBP variability is based on observational results obtained from our experimental data and might be validated by RBP engineering methods, such as structure-guided mutagenesis or RBP swapping (52). Curiously, it is worth mentioning that RBP versatility seems to be associated with a given protein structure. The most versatile and variable RBPs were long chaperone-dependent tail spikes with a triple helix Dpo domain, which had a similar distribution of mutations. In contrast, most specific RBPs were tail spike proteins with a right-handed β-helix Dpo domain.

**Figure 4.**
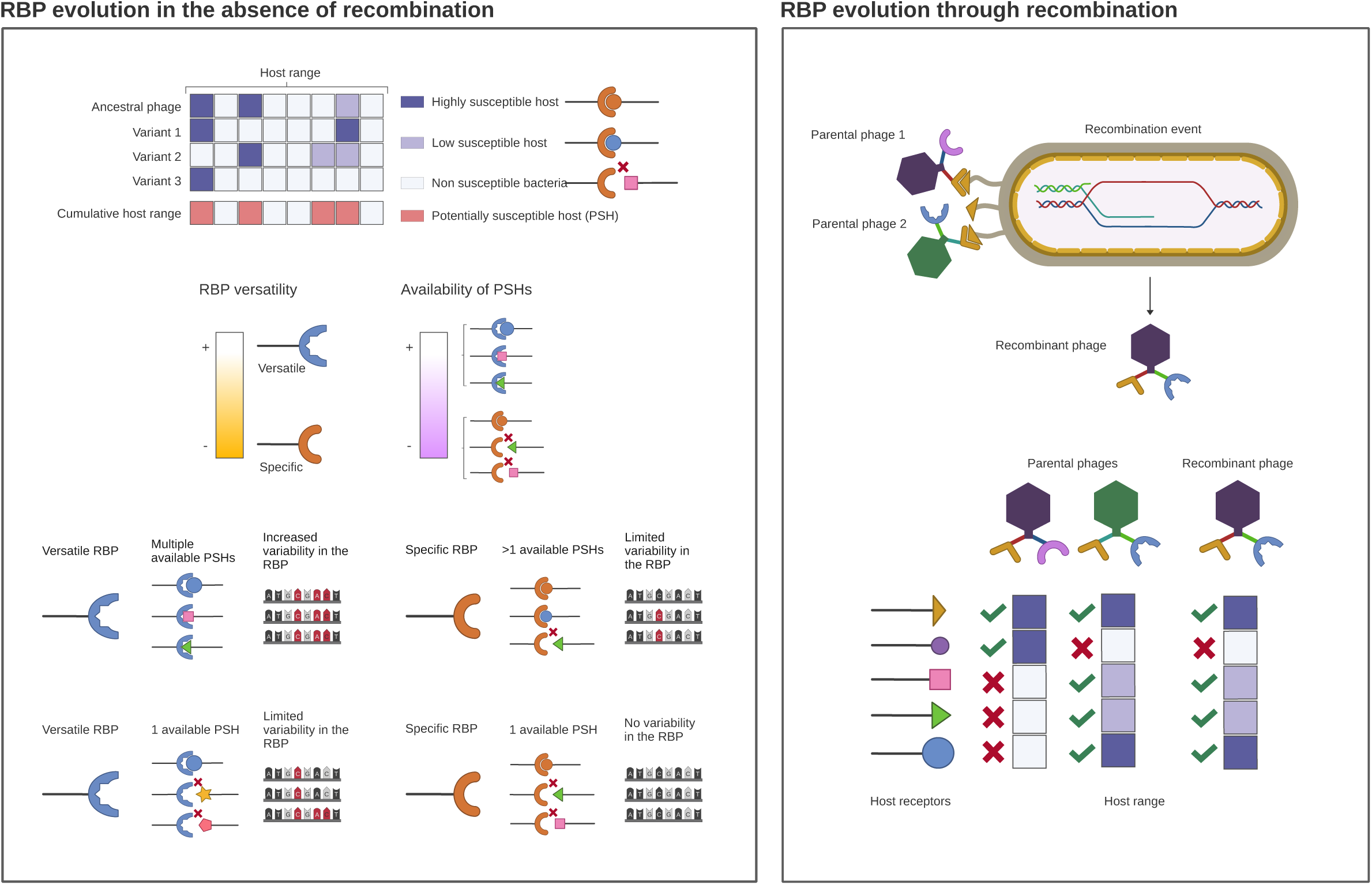
Proposed model for host range evolution based on receptor binding proteins variability. Based on experimental data obtained in this work, we propose a model distinguishing two main scenarios, in the absence and presence of recombination. In the first one, the host range of each variant includes differentially susceptible hosts. The cumulative host range is the sum of the host ranges of the ancestral phage and its variants, including all the potentially susceptible hosts (PSHs). A receptor binding protein (RBP) is considered more versatile or more specific regarding the number of receptors it can target. The degree of versatility of an RBP together with the availability of PSHs in a given environment determines the accumulation of mutations over time. In the second scenario, recombination events can affect versatile or specific RBPs, rapidly modifying the host range.

During evolution, mutations in other genes were accumulated in the lineages. Given the huge arsenal of anti-phage defense systems that bacteria encode to protect from phage infection, it is probable that some other mutations have emerged by escaping these systems. Mutation in targets of nucleic acid degradation systems is a typical escape strategy of phages (19,53). Moreover, mutations in proteins that act as triggers for defense systems can allow phage escape. This includes structural proteins, involved in replication, or direct inhibitors of host defenses (19,54). The central tail fiber J is an interesting example. Amino acid changes in the C-terminal portion of this protein observed in our experiment have been previously associated with a counteradaptation to the Tai immune system. In addition, this protein can be involved in the recognition and binding process (55). Other interesting mutations detected are the large deletions in specific phages, removing non-essential genes and improving replication (56–58), as observed here.

This work represents a significant advancement in our understanding of the interactions between bacteria and their phages, with important implications in both industrial and clinical settings, especially in an era of rising antibiotic resistance. Knowing the basis of host range evolution and phage dynamics in host-diverse environments is relevant to predicting how the phages, used in single phage preparations or cocktails, would impact microbial communities, such as the human microbiota. In summary, our work contributes to elucidating the molecular mechanisms underlying the evolution of the host range of *Klebsiella* phages and a better understanding of phage interactions with other phages and hosts in a host-diverse community.

## Materials and methods

### Bacterial strains

The *Klebsiella* reference strains collection, corresponding to the 77 *Klebsiella* reference serotypes, was purchased from the Statens Serum Institut (Copenhagen, Denmark). Genomes of 62 of the 77 reference strains were available online (Supplementary Table 2). The collection includes diverse *Klebsiella* species: *K. pneumoniae*, *K. planticola*, *K. oxytoca*, *K. ozaenae*, and *K. terrigena*. All bacteria were grown in Luria-Bertani (LB) with CaCl2 (3.78mM) broth at 37°C at 180 rpm.

### Bacterial pool for the experimental evolution

A bacterial pool encompassing 39 reference strains of 4 different *Klebsiella* species (39 different K-types) was created for the experimental evolution, as a model for a host-diverse environment (Supplementary Table 2). An exponential culture of each bacterial strain was prepared at an OD620 ≈ 0.2 (∼10^8^ colony forming units (CFU)/mL). All cultures were mixed in the same proportion to elaborate the stock of the bacterial population for the passages. The combined culture was concentrated and stored at -70°C in aliquots with 20% of glycerol for its preservation.

### Phage cocktail preparation

The 12-phage cocktail used for the experimental evolution was previously designed and tested in an in-house collection of *K. pneumoniae* clinical strains (27). This cocktail combined large dsDNA specialist and generalist phages, with a continuum of host ranges and genomically diverse, including representatives of 9 different genera (2 *Sugarlandvirus*, 2 *Przondovirus*, 2 *Mydovirus*, 1 *Vectrevirus*, 1 *Taipeivirus*, 1 *Jiaodavirus*, 1 *Drexlerviridae* unclassified, 1 unclassified family) (Supplementary Table 1). For the cocktail elaboration, phages were propagated separately in a final volume of 5mL LB+CaCl2 broth using its isolation strain. Phages were finally combined in the cocktail to a final titer of 10^8^ PFU/mL per phage. Assemblies of the ancestral phages were already available (27). Genomes were reassembled using Unicycler (version 0.5.0) and annotated using Pharokka (v1.4.1). Specific identification of RBPs was performed by protein alignment with the already available annotation of the ancestral genomes, where RBP annotation was specifically refined.

### Experimental evolution of the phage cocktail in a host-diverse environment

A bacterial pool aliquot was resuspended in LB (OD620 ≈ 0.2). For the first inoculum (passage 0), 10^8^ PFU/mL per phage of the initial cocktail were added to the bacterial pool in three different tubes in a final titer of 10^6^ plaque forming units (PFU)/mL per phage, initiating three lineages of evolution. Tubes for each lineage were incubated for 3.5 hours at 37°C at 750 rpm. After that, they were centrifuged to eliminate bacteria (passage 1) and the supernatant was diluted to adjust the inoculum for the next passage. To maintain an adjusted titer during the experiment, every 2 passages the phage population was titrated in at least 3 different strains. A total of 69 passages were performed for the three independent lineages.

### Experimental evolution of single phages

Single phages were evolved in parallel to validate the results from the experimental evolution in a host-diverse environment. Phage initial inoculum was 10^8^ PFU/mL for each phage. The evolution was performed similarly in three independent lineages per phage during 20 passages, as described above.

### Host range evaluation

For the phage cocktail, the host range was firstly tested for passage 0 (ancestral phage cocktail at 10^8^ PFU/mL per phage), to detect possible phage-host interactions not reported at the conditions assayed in our previous work (27). In addition, the host range was evaluated throughout the experimental evolution, in passages 20, 40, and 69. Serial dilutions for each lineage (and passage) were tested for the phage populations using the spot test technique (dilutions from 1 to 10^-6^) in the 77 reference collection as described (27). An interaction was considered positive only if single plaques were consistently observed in the serial dilutions in the different replicates. Turbid spots only present in the 10^-1^ dilution were considered ambiguous interactions. The absence of a visual spot was considered a negative interaction. Two replicates of the experiment were performed, and three for the doubtful cases. This was similarly performed for passage 20 of single-phage evolution lineages.

### Single-plaque isolation of phages with new host rang

Plaque isolation was performed in newly adapted hosts according to the infectivity matrix previously published (27) and in 4 already susceptible hosts. A dilution of the passage corresponding to the limiting dilution observed in the host range evaluation for each strain was plated, having a total of 96 strain- passage combinations. When plaques were observed, they were recovered and purified. After phage purification, they were propagated in its isolation strain and concentrated using the Concentrating Pipette Select System (Innovaprep) with pipettes Ultra (<0.05 µm). The host range of the single-plaque phages was evaluated by spot test as described previously.

### Viral genome sequencing

Passages 0, 40, and 69 were sequenced from the cocktail evolution, as single plaques, and passages 0 and 20 from single-phage evolutions. Removal of host DNA and digestion of phage capsids were performed as described before (26). Extraction and purification of DNA were done using DNA Clean & Concentrator 5-Kit (Zymo) for the passages and using Maxwell PureFood GMO and Authentication Kit (Promega) with Maxwell RSC Instrument (Promega) for single plaques.

Sequencing libraries were prepared using the Illumina Nextera XT DNA kit (paired-end reads 2x250 bp). Reads were generated in the Illumina MiSeq platform with MiSeq Reagent Kit v2 for the passages of the cocktail evolution, with MiSeq Reagent Kit v3 (2x250 bp) for the passages of single-phage evolutions, and with MiSeq Reagent Kit v2 nano for the single plaques. Sequencing read quality was assessed using FastQC software (version 0.11.9, Babraham Bioinformatics) (59). De novo genome assembly for the passages was carried out with the ‘metaspades’ function of SPAdes (version 3.15.4) (60,61). For genomic data from single plaques, Unicycler (version 0.5.0) (62) in combination with SPAdes (version 3.15.4) was performed. If necessary, a subset of 10.000 reads was used for the assembly and refined using pilon (version 1.24) (63). Read mapping for coverage calculation was performed using BBMap (64).

### Genomic characterization of the phage populations

Intergenomic similarity of phages was assessed using VIRIDIC (65). To avoid nonspecific read mapping, only phages with an intergenomic similarity lower than 2 with any other phage in the cocktail were considered sufficiently different to be analyzed by variant calling. For this reason, small regions with a high percentage of identity (higher than 80%) were excluded from the analysis. For phages with higher intergenomic similarities but between which there is sufficient difference in sequencing depth (ratio of the percentage of reads per position lower than 20:80), variant calling analysis was also performed for the phage with the highest sequencing depth. Reads were mapped using BWA (version 0.7.17) (66) and the variants were called using LoFreq (version 2.1.5) (67). In parallel, read mapping was visualized using Integrative Genomics Viewer (IGV) (68). CDS with mutations were translated using MEGAX (69) to check whether the mutations were synonyms or non-synonyms, and amino acid changes. For visualization of mutations detected along the genome of the ancestral phage, the mutation rate was calculated on sliding windows of 500 bp for each lineage. A graphic representation of this value was overlaid with the graphic representation of ancestral phage annotation obtained with the R package gggenomes (70).

To analyze the possible activation of prophages from the bacterial population, unmapped reads were assembled using the ‘metaspades’ function of SPAdes (60,61). Viral contigs were selected using VirSorter2 (version 2.2.4, only including dsDNAphages with a minimum length of 2000 bp) (71) and CheckV (version 1.0.1, database v1.5) (72). Contigs encoding less than two genes or with any viral gene detected were discarded for further analyses. The selected contigs were aligned with available bacterial genomes of the strains included in the bacterial pool using BLAST (73). Those with a similarity up to 99% in at least 99% of their length with an E-value = 0.0 were considered positive hits. Hits were confirmed by read mapping against corresponding bacterial strains and visualized with IGV. Prophage activation was assumed if reads were uniquely mapped in a specific region. Bacteria with positive hits were searched for prophages in their genomes using PHASTEST (74), and viral contigs that hit with each bacteria were compared using BLAST (73) with prophages detected to confirm their proviral origin.

Analysis of single-phage evolution populations was done similarly. Reads obtained in each sample were assembled with SPAdes. Contigs corresponding to the evolved phage were identified with BLAST. Other viral contigs were selected and analyzed as described before with VirSorter2 and CheckV. Variant calling was performed as described previously. Completely recovered active prophages were aligned with available bacterial genomes with BLAST to verify their origin, and with contigs from other evolution lineages to find the fragmented genome.

### Analysis of variability of the evolved phages

Genomes of the evolved phage variants obtained from single plaque isolation were classified and reordered based on their parentals. Functional annotation was carried out using Pharokka (v1.4.1) (75). Genome alignment was performed with MAFFT (version 7.520) (76) to evaluate variable positions. For the recombinant phages isolated, recombination events were detected using RDP4 (77). Recombination events were considered when confirmed by at least 5 of the 7 utilized methods: RDP, GENECONV, MaxChi, BootScan, Chimaera, 3Seq, and SiScan. Each evolved variant was named related to its ancestral phage, including “_evo” (g.e. K50_evo1 as a non-recombinant evolved variant of phage K50PH164C1). In the case of the recombinants, the major parental is used adding “_rec” (g.e. K7_rec1 as a recombinant whose major parent is phage K7PH164C4).

We focused on analyzing the RBPs and the central tail fiber J (if present) of the evolved phages. Amino acid sequences of these proteins were aligned using MAFFT (76) to find variable positions in the amino acid sequence. Protein domains were identified when possible using InterProScan (78) or manually (27) based on the 3D structure of the protein, predicted with AlphaFold Server, which uses AlphaFold3 (79). The coloring of regions of interest and representation of variable positions of the protein sequence alignment as ‘spheres’ was done with Pymol (80). HHpred (81) was used to assess similarity to other protein groups.

## Acknowledgements

We thank Amanda Martínez and Sandra Albert for technical assistance. This research was funded by project PID2020-112835RA-I00 funded by MCIN/AEI/10.13039/501100011033, and project SEJIGENT/2021/014 funded by Conselleria d’Innovació, Universitats, Ciència i Societat Digital (Generalitat Valenciana) to P.D.-C. C.F.-G. was funded by a PhD fellowship Atracció de Talent UV-INV_PREDOC-1913324 from Universitat de València. P.D.-C. was financially supported by a Ramón y Cajal contract RYC2019-028015-I funded by MCIN/AEI/10.13039/501100011033, ESF Invest in your future.

## Author contributions

C.F.-G. performed the experiments, the computational analyses, data analysis, visualization, and manuscript writing. P.D.-C. provided reagents, conceived the project, designed the research, contributed to data analysis, revised the manuscript, and conducted the supervision. All authors read and approved the final manuscript.

## References

1. Zhang YZ, Shi M, Holmes EC. Using Metagenomics to Characterize an Expanding Virosphere. Cell. 2018 Mar 8;172(6):1168–72.

2. Koskella B, Hernandez CA, Wheatley RM. Understanding the Impacts of Bacteriophage Viruses: From Laboratory Evolution to Natural Ecosystems. Annu Rev Virol. 2022 Sep 29;9(Volume 9, 2022):57–78.

3. Manrubia SC, Lázaro E. Viral evolution. Phys Life Rev. 2006 Jun 1;3(2):65–92.

4. Kawecki TJ, Lenski RE, Ebert D, Hollis B, Olivieri I, Whitlock MC. Experimental evolution. Trends Ecol Evol. 2012 Oct 1;27(10):547–60.

5. Clokie MR, Millard AD, Letarov AV, Heaphy S. Phages in nature. Bacteriophage. 2011 Jan;1(1):31–45.

6. Jonge PA de, Nobrega FL, Brouns SJJ, Dutilh BE. Molecular and Evolutionary Determinants of Bacteriophage Host Range. Trends Microbiol. 2019 Jan 1;27(1):51–63.

7. Yu P, Mathieu J, Li M, Dai Z, Alvarez PJJ. Isolation of Polyvalent Bacteriophages by Sequential Multiple-Host Approaches. Appl Environ Microbiol. 2016 Feb 1;82(3):808–15.

8. Sant DG, Woods LC, Barr JJ, McDonald MJ. Host diversity slows bacteriophage adaptation by selecting generalists over specialists. Nat Ecol Evol. 2021 Mar;5(3):350–9.

9. Bono LM, Gensel CL, Pfennig DW, Burch CL. Evolutionary rescue and the coexistence of generalist and specialist competitors: an experimental test. Proc R Soc B Biol Sci. 2015 Dec 22;282(1821):20151932.

10. Ouyang R, Ongenae V, Muok A, Claessen D, Briegel A. Phage fibers and spikes: a nanoscale Swiss army knife for host infection. Curr Opin Microbiol. 2024 Feb;77:102429.

11. Holtzman T, Globus R, Molshanski-Mor S, Ben-Shem A, Yosef I, Qimron U. A continuous evolution system for contracting the host range of bacteriophage T7. Sci Rep. 2020 Jan 15;10(1):307.

12. Pas C, Latka A, Fieseler L, Briers Y. Phage tailspike modularity and horizontal gene transfer reveals specificity towards E. coli O-antigen serogroups. Virol J. 2023 Aug 7;20(1):174.

13. Oliveira PH, Touchon M, Rocha EPC. The interplay of restriction-modification systems with mobile genetic elements and their prokaryotic hosts. Nucleic Acids Res. 2014;42(16):10618–31.

14. Chopin MC, Chopin A, Bidnenko E. Phage abortive infection in lactococci: variations on a theme. Curr Opin Microbiol. 2005 Aug;8(4):473–9.

15. Makarova KS, Wolf YI, Alkhnbashi OS, Costa F, Shah SA, Saunders SJ, et al. An updated evolutionary classification of CRISPR-Cas systems. Nat Rev Microbiol. 2015 Nov;13(11):722–36.

16. Bernheim A, Bikard D, Touchon M, Rocha EPC. Atypical organizations and epistatic interactions of CRISPRs and cas clusters in genomes and their mobile genetic elements. Nucleic Acids Res. 2020 Jan 24;48(2):748–60.

17. Doron S, Melamed S, Ofir G, Leavitt A, Lopatina A, Keren M, et al. Systematic discovery of antiphage defense systems in the microbial pangenome. Science. 2018 Mar 2;359(6379):eaar4120.

18. Tesson F, Hervé A, Mordret E, Touchon M, d’Humières C, Cury J, et al. Systematic and quantitative view of the antiviral arsenal of prokaryotes. Nat Commun. 2022 May 10;13(1):2561.

19. Mayo-Muñoz D, Pinilla-Redondo R, Camara-Wilpert S, Birkholz N, Fineran PC. Inhibitors of bacterial immune systems: discovery, mechanisms and applications. Nat Rev Genet. 2024 Apr;25(4):237–54.

20. Scholl D, Adhya S, Merril C. Escherichia coli K1’s capsule is a barrier to bacteriophage T7. Appl Environ Microbiol. 2005 Aug;71(8):4872–4.

21. Rendueles O, Garcia-Garcerà M, Néron B, Touchon M, Rocha EPC. Abundance and co-occurrence of extracellular capsules increase environmental breadth: Implications for the emergence of pathogens. PLoS Pathog. 2017 Jul;13(7):e1006525.

22. Mostowy RJ, Holt KE. Diversity-Generating Machines: Genetics of Bacterial Sugar-Coating. Trends Microbiol. 2018 Dec;26(12):1008–21.

23. Latka A, Lemire S, Grimon D, Dams D, Maciejewska B, Lu T, et al. Engineering the Modular Receptor-Binding Proteins of Klebsiella Phages Switches Their Capsule Serotype Specificity. mBio. 2021 May 4;12(3):10.1128/mbio.00455-21.

24. Squeglia F, Maciejewska B, Łątka A, Ruggiero A, Briers Y, Drulis-Kawa Z, et al. Structural and Functional Studies of a *Klebsiella* Phage Capsule Depolymerase Tailspike: Mechanistic Insights into Capsular Degradation. Structure. 2020 Jun 2;28(6):613–624.e4.

25. Popova AV, Shneider MM, Arbatsky NP, Kasimova AA, Senchenkova SN, Shashkov AS, et al. Specific Interaction of Novel Friunavirus Phages Encoding Tailspike Depolymerases with Corresponding Acinetobacter baumannii Capsular Types. J Virol. 2021 Mar 1;95(5):e01714–20, JVI.01714-20.

26. Beamud B, García-González N, Gómez-Ortega M, González-Candelas F, Domingo-Calap P, Sanjuan R. Genetic determinants of host tropism in Klebsiella phages. Cell Rep. 2023 Feb 28;42(2):112048.

27. Ferriol-González C, Concha-Eloko R, Bernabéu-Gimeno M, Fernández-Cuenca F, Cañada-García JE, García-Cobos S, et al. Targeted phage hunting to specific Klebsiella pneumoniae clinical isolates is an efficient antibiotic resistance and infection control strategy. Microbiol Spectr. 2024 Aug 28;0(0):e00254–24.

28. Thorpe HA, Booton R, Kallonen T, Gibbon MJ, Couto N, Passet V, et al. A large-scale genomic snapshot of Klebsiella spp. isolates in Northern Italy reveals limited transmission between clinical and non-clinical settings. Nat Microbiol. 2022 Dec;7(12):2054–67.

29. Martin RM, Bachman MA. Colonization, Infection, and the Accessory Genome of Klebsiella pneumoniae. Front Cell Infect Microbiol. 2018;8:4.

30. Lau HY, Huffnagle GB, Moore TA. Host and microbiota factors that control Klebsiella pneumoniae mucosal colonization in mice. Microbes Infect. 2008 Oct;10(12–13):1283– 90.

31. David S, Reuter S, Harris SR, Glasner C, Feltwell T, Argimon S, et al. Epidemic of carbapenem-resistant Klebsiella pneumoniae in Europe is driven by nosocomial spread. Nat Microbiol. 2019 Nov;4(11):1919–29.

32. European Antimicrobial Resistance Collaborators. The burden of bacterial antimicrobial resistance in the WHO European region in 2019: a cross-country systematic analysis. Lancet Public Health. 2022 Nov;7(11):e897–913.

33. Mak ANS, Lambert AR, Stoddard BL. Folding, DNA Recognition, and Function of GIY-YIG Endonucleases: Crystal Structures of R.Eco29kI. Structure. 2010 Oct 13;18(10):1321–31.

34. Concha-Eloko R, Barberán-Martínez P, Sanjuán R, Domingo-Calap P. Broad-range capsule-dependent lytic Sugarlandvirus against Klebsiella sp. Microbiol Spectr. 2023 Oct 26;11(6):e04298–22.

35. Murphy J, Mahony J, Ainsworth S, Nauta A, van Sinderen D. Bacteriophage Orphan DNA Methyltransferases: Insights from Their Bacterial Origin, Function, and Occurrence. Appl Environ Microbiol. 2013 Dec 15;79(24):7547–55.

36. Tal N, Millman A, Stokar-Avihail A, Fedorenko T, Leavitt A, Melamed S, et al. Bacteria deplete deoxynucleotides to defend against bacteriophage infection. Nat Microbiol. 2022 Aug;7(8):1200–9.

37. Klemm BP, Singh D, Smith CE, Hsu AL, Dillard LB, Krahn JM, et al. Mechanism by which T7 bacteriophage protein Gp1.2 inhibits Escherichia coli dGTPase. Proc Natl Acad Sci. 2022 Sep 13;119(37):e2123092119.

38. Jakob N, Hammerl JA, Swierczewski BE, Würstle S, Bugert JJ. Appelmans Protocol for in vitro Klebsiella pneumoniae phage host range expansion leads to induction of a novel temperate linear plasmid prophage vB_KpnS-KpLi5. bioRxiv. 2023 Aug 6;2023–08.

39. Peters TL, Schow J, Spencer E, Leuven JV, Wichman H, Miller C. Directed evolution of bacteriophages: impacts of prolific prophage. bioRxiv. 2024 Jun 29;2024–06.

40. Tisza MJ, Buck CB. A catalog of tens of thousands of viruses from human metagenomes reveals hidden associations with chronic diseases. Proc Natl Acad Sci U S A. 2021 Jun 8;118(23):e2023202118.

41. Di Pilato V, Aiezza N, Viaggi V, Antonelli A, Principe L, Giani T, et al. KPC-53, a KPC-3 Variant of Clinical Origin Associated with Reduced Susceptibility to Ceftazidime-Avibactam. Antimicrob Agents Chemother. 2020 Dec 16;65(1):10.1128/aac.01429-20.

42. Gorodnichev RB, Volozhantsev NV, Krasilnikova VM, Bodoev IN, Kornienko MA, Kuptsov NS, et al. Novel Klebsiella pneumoniae K23-Specific Bacteriophages From Different Families: Similarity of Depolymerases and Their Therapeutic Potential. Front Microbiol. 2021;12:669618.

43. Gorrie CL, Mirčeta M, Wick RR, Judd LM, Lam MMC, Gomi R, et al. Genomic dissection of Klebsiella pneumoniae infections in hospital patients reveals insights into an opportunistic pathogen. Nat Commun. 2022 May 31;13(1):3017.

44. Bull JJ, Wichman HA, Krone SM. Modeling the Directed Evolution of Broad Host Range Phages. Antibiotics. 2022 Nov 27;11(12):1709.

45. Hibbing ME, Fuqua C, Parsek MR, Peterson SB. Bacterial competition: surviving and thriving in the microbial jungle. Nat Rev Microbiol. 2010 Jan;8(1):15–25.

46. Bailey ZM, Igler C, Wendling CC. Prophage maintenance is determined by environment-dependent selective sweeps rather than mutational availability. Curr Biol. 2024 Apr 22;34(8):1739–1749.e7.

47. Ford BE, Sun B, Carpino J, Chapler ES, Ching J, Choi Y, et al. Frequency and Fitness Consequences of Bacteriophage Φ6 Host Range Mutations. PLoS ONE. 2014 Nov 19;9(11):e113078.

48. Burmeister AR, Tzintzun-Tapia E, Roush C, Mangal I, Barahman R, Bjornson RD, et al. Experimental Evolution of the TolC-Receptor Phage U136B Functionally Identifies a Tail Fiber Protein Involved in Adsorption through Strong Parallel Adaptation. Appl Environ Microbiol. 2023 May 16;89(6):e00079–23.

49. Akusobi C, Chan BK, Williams ESCP, Wertz JE, Turner PE. Parallel Evolution of Host-Attachment Proteins in Phage PP01 Populations Adapting to Escherichia coli O157:H7. Pharmaceuticals. 2018 Jun;11(2):60.

50. Pan YJ, Lin TL, Chen CC, Tsai YT, Cheng YH, Chen YY, et al. Klebsiella Phage ΦK64-1 Encodes Multiple Depolymerases for Multiple Host Capsular Types. J Virol. 2017 Feb 28;91(6):10.1128/jvi.02457-16.

51. Zhou Y, Li L, Han K, Wang L, Cao Y, Ma D, et al. A Polyvalent Broad-Spectrum Escherichia Phage Tequatrovirus EP01 Capable of Controlling Salmonella and Escherichia coli Contamination in Foods. Viruses. 2022 Jan 29;14(2):286.

52. Dunne M, Prokhorov NS, Loessner MJ, Leiman PG. Reprogramming bacteriophage host range: design principles and strategies for engineering receptor binding proteins. Curr Opin Biotechnol. 2021 Apr 1;68:272–81.

53. Pleška M, Guet CC. Effects of mutations in phage restriction sites during escape from restriction-modification. Biol Lett. 2017 Dec;13(12):20170646.

54. Stokar-Avihail A, Fedorenko T, Hör J, Garb J, Leavitt A, Millman A, et al. Discovery of phage determinants that confer sensitivity to bacterial immune systems. Cell. 2023 Apr 27;186(9):1863–1876.e16.

55. He L, Miguel-Romero L, Patkowski JB, Alqurainy N, Rocha EPC, Costa TRD, et al. Tail assembly interference is a common strategy in bacterial antiviral defenses. Nat Commun. 2024 Aug 30;15(1):7539.

56. Tom EF, Molineux IJ, Paff ML, Bull JJ. Experimental evolution of UV resistance in a phage. PeerJ. 2018 Jul 9;6:e5190.

57. Edwards KF, Steward GF, Schvarcz CR. Making sense of virus size and the tradeoffs shaping viral fitness. Ecol Lett. 2021;24(2):363–73.

58. Yuan S, Shi J, Jiang J, Ma Y. Genome-scale top-down strategy to generate viable genome-reduced phages. Nucleic Acids Res. 2022 Dec 9;50(22):13183–97.

59. Andrews S. FastQC: a quality control tool for high throughput sequence data. 2010.

60. Bankevich A, Nurk S, Antipov D, Gurevich AA, Dvorkin M, Kulikov AS, et al. SPAdes: A New Genome Assembly Algorithm and Its Applications to Single-Cell Sequencing. J Comput Biol. 2012 May;19(5):455–77.

61. Nurk S, Meleshko D, Korobeynikov A, Pevzner PA. metaSPAdes: a new versatile metagenomic assembler. Genome Res. 2017 Jan 5;27(5):824–34.

62. Wick RR, Judd LM, Gorrie CL, Holt KE. Unicycler: Resolving bacterial genome assemblies from short and long sequencing reads. PLoS Comput Biol. 2017 Jun;13(6):e1005595.

63. Walker BJ, Abeel T, Shea T, Priest M, Abouelliel A, Sakthikumar S, et al. Pilon: An Integrated Tool for Comprehensive Microbial Variant Detection and Genome Assembly Improvement. PLOS ONE. 2014 Nov 19;9(11):e112963.

64. Bushnell B. BBMap: A Fast, Accurate, Splice-Aware Aligner. 2014 Mar 19 [cited 2024 Sep 3]; Available from: https://escholarship.org/uc/item/1h3515gn

65. Moraru C, Varsani A, Kropinski AM. VIRIDIC-A Novel Tool to Calculate the Intergenomic Similarities of Prokaryote-Infecting Viruses. Viruses. 2020 Nov 6;12(11):1268.

66. Li H, Durbin R. Fast and accurate short read alignment with Burrows-Wheeler transform. Bioinforma Oxf Engl. 2009 Jul 15;25(14):1754–60.

67. Wilm A, Aw PPK, Bertrand D, Yeo GHT, Ong SH, Wong CH, et al. LoFreq: a sequence-quality aware, ultra-sensitive variant caller for uncovering cell-population heterogeneity from high-throughput sequencing datasets. Nucleic Acids Res. 2012 Dec;40(22):11189– 201.

68. Robinson JT, Thorvaldsdottir H, Turner D, Mesirov JP. igv.js: an embeddable JavaScript implementation of the Integrative Genomics Viewer (IGV). Bioinformatics. 2023 Jan 1;39(1):btac830.

69. Kumar S, Stecher G, Li M, Knyaz C, Tamura K. MEGA X: Molecular Evolutionary Genetics Analysis across Computing Platforms. Mol Biol Evol. 2018 Jun 1;35(6):1547–9.

70. Hackl T, Ankenbrand M, van Adrichem B. gggenomes: A Grammar of Graphics for Comparative Genomics [Internet]. 2024. Available from: https://github.com/thackl/gggenomes

71. Guo J, Bolduc B, Zayed AA, Varsani A, Dominguez-Huerta G, Delmont TO, et al. VirSorter2: a multi-classifier, expert-guided approach to detect diverse DNA and RNA viruses. Microbiome. 2021 Feb 1;9(1):37.

72. Nayfach S, Camargo AP, Schulz F, Eloe-Fadrosh E, Roux S, Kyrpides NC. CheckV assesses the quality and completeness of metagenome-assembled viral genomes. Nat Biotechnol. 2021 May;39(5):578–85.

73. Altschul SF, Gish W, Miller W, Myers EW, Lipman DJ. Basic local alignment search tool. J Mol Biol. 1990 Oct 5;215(3):403–10.

74. Wishart DS, Han S, Saha S, Oler E, Peters H, Grant JR, et al. PHASTEST: faster than PHASTER, better than PHAST. Nucleic Acids Res. 2023 Jul 5;51(W1):W443–50.

75. Bouras G, Nepal R, Houtak G, Psaltis AJ, Wormald PJ, Vreugde S. Pharokka: a fast scalable bacteriophage annotation tool. Bioinforma Oxf Engl. 2023 Jan 1;39(1):btac776.

76. Katoh K, Misawa K, Kuma K ichi, Miyata T. MAFFT: a novel method for rapid multiple sequence alignment based on fast Fourier transform. Nucleic Acids Res. 2002 Jul 15;30(14):3059–66.

77. Martin DP, Murrell B, Golden M, Khoosal A, Muhire B. RDP4: Detection and analysis of recombination patterns in virus genomes. Virus Evol. 2015;1(1):vev003.

78. Jones P, Binns D, Chang HY, Fraser M, Li W, McAnulla C, et al. InterProScan 5: genome-scale protein function classification. Bioinformatics. 2014 May 1;30(9):1236–40.

79. Abramson, J., Adler, J., Dunger, J. et al. Accurate structure prediction of biomolecular interactions with AlphaFold 3. Nature. 2024 Jun. 630, 493–500.

80. Schrödinger L. The PyMOL Molecular Graphics System. 2020.

81. Söding J, Biegert A, Lupas AN. The HHpred interactive server for protein homology detection and structure prediction. Nucleic Acids Res. 2005 Jul 1;33:W244–8.

