## Supplementary Figures for "Evolutionary strategies of *Klebsiella* phages in a host-diverse environment"

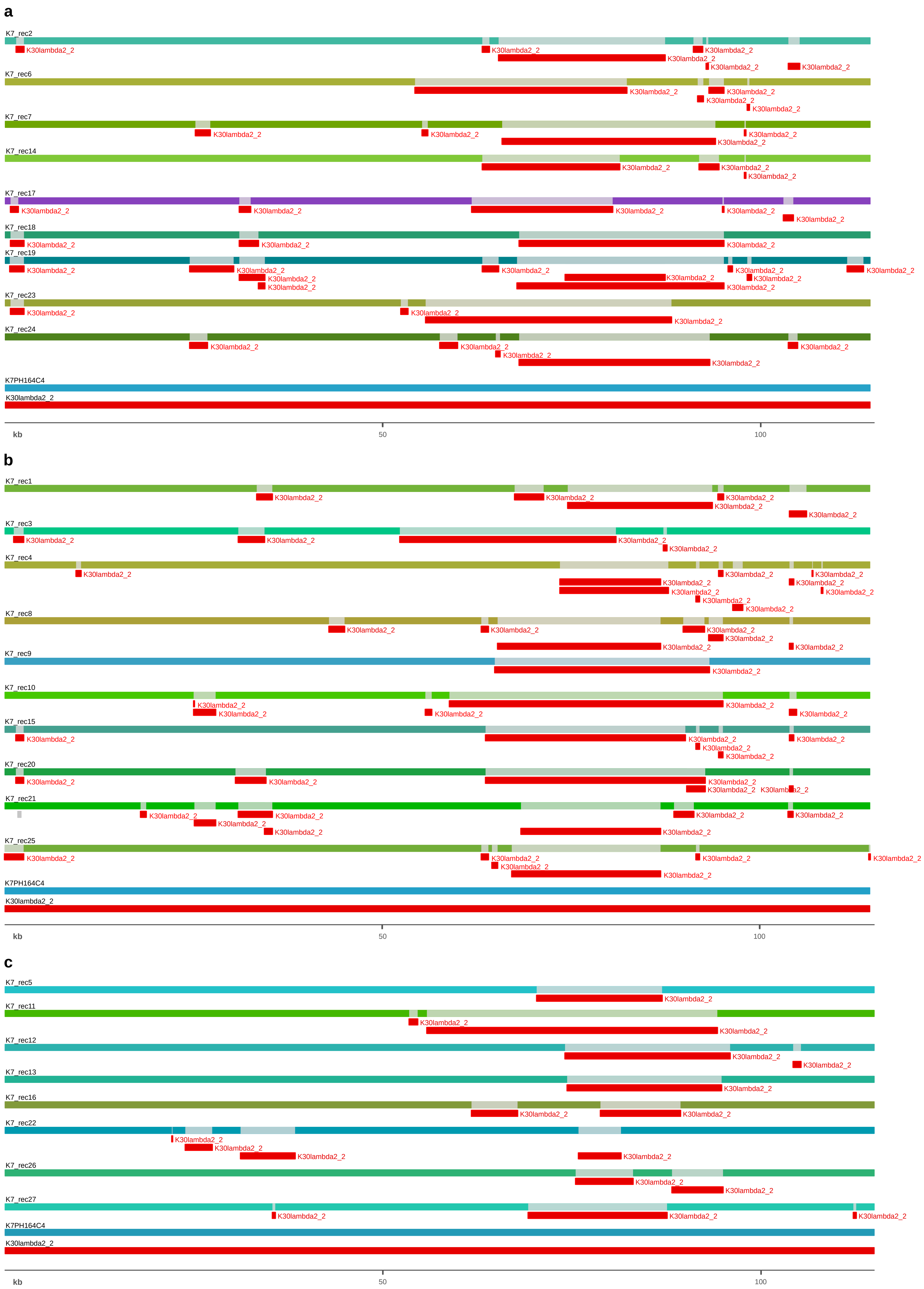
a
K7_rec2
K7_rec6
K7_rec7
K7_rec14
K7_rec17
K7_rec18
K7_rec19
K7_rec23
K7_rec24
K7PH164C4
K30lambda2_2
kb
50
100
b
K7_rec1
K7_rec3
K7_rec4
K7_rec8
K7_rec9
K7_rec10
K7_rec15
K7_rec20
K7_rec21
K7_rec25
K7PH164C4
K30lambda2_2
kb
50
100
c
K7_rec5
K7_rec11
K7_rec12
K7_rec13
K7_rec16
K7_rec22
K7_rec26
K7_rec27
K7PH164C4
K30lambda2_2
kb
50
100

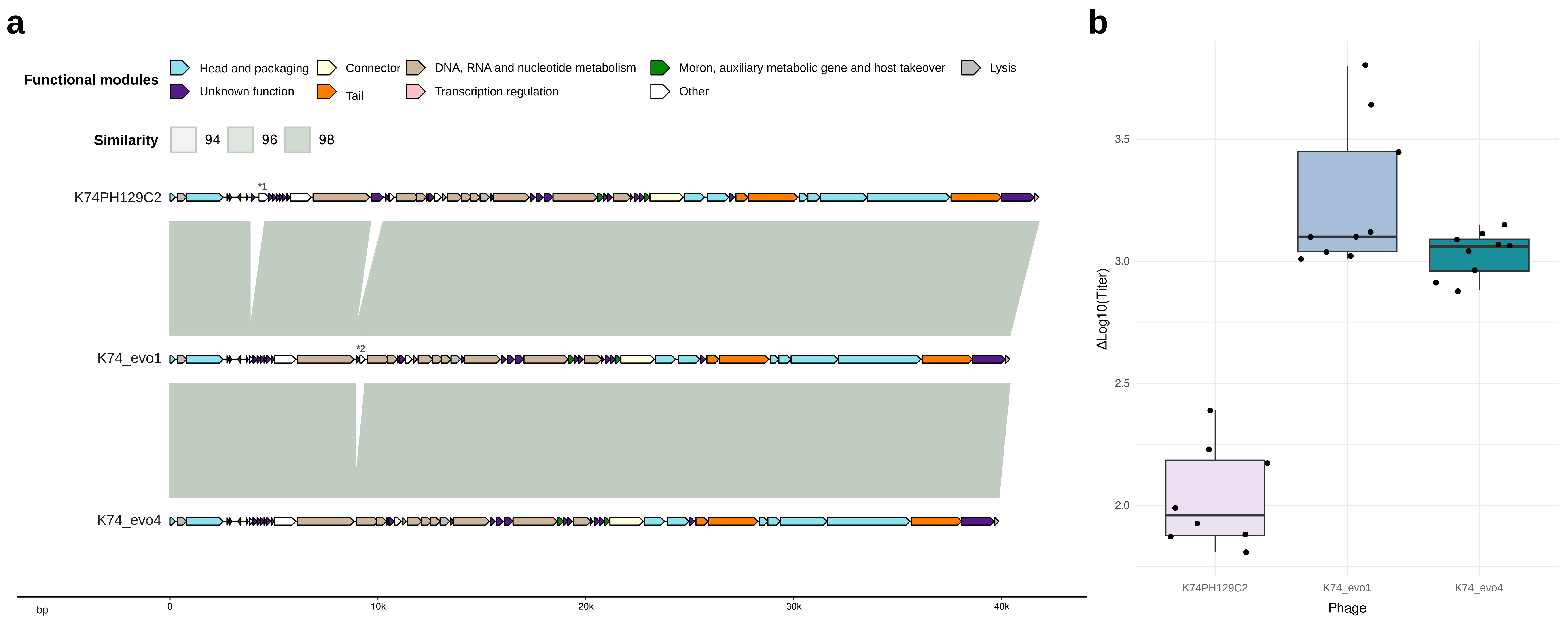

a
Functional modules
Head and packaging
Unknown function
Connector
Tail
DNA, RNA and nucleotide metabolism
Transcription regulation
Moron, auxiliary metabolic gene and host takeover
Other
Lysis
Similarity
94
96
98
K74PH129C2
*1
K74_evo1
*2
K74_evo4
bp
0
10k
20k
30k
40k
b
ΔLog10(Titer)
3.5
3.0
2.5
2.0
K74PH129C2
K74_evo1
K74_evo4
Phage

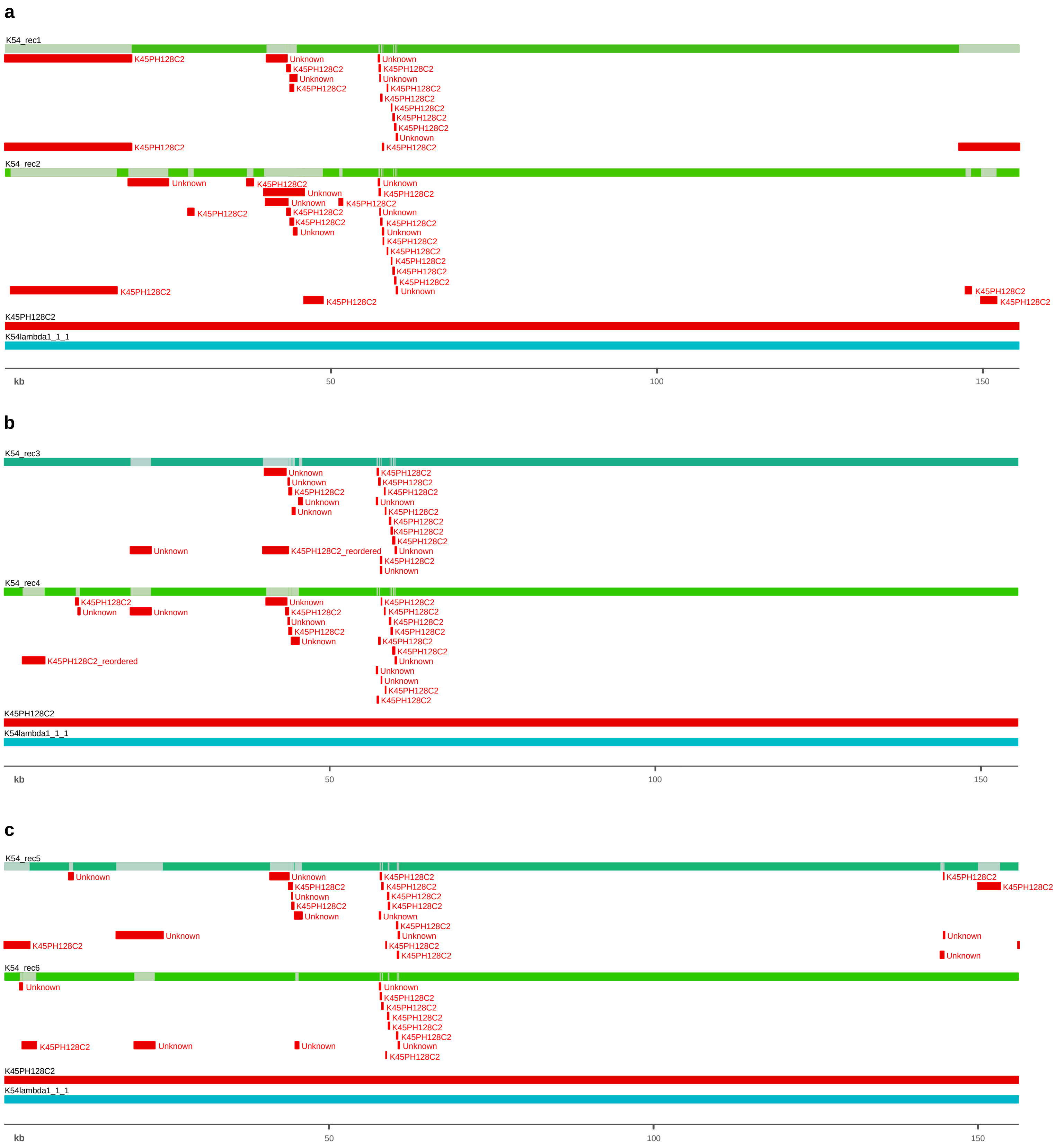
a
K54_rec1
K54_rec2
K45PH128C2
K54lambda1_1_1
Unknown
K45PH128C2
kb
50
100
150
b
K54_rec3
K54_rec4
K45PH128C2_reordered
K45PH128C2
K54lambda1_1_1
Unknown
kb
50
100
150
c
K54_rec5
K54_rec6
K45PH128C2
K54lambda1_1_1
Unknown
kb
50
100
150

**Supplementary Figures**

Supplementary Figure 1. **Graphic representation of recombinant fragments of evolved *Sugarlandviruses*.** The figure illustrates the whole genome alignment of all recombinants of each lineage with parental phages. The genome size bar corresponds to positions in the alignment. K7PH164C4 is the major parental and its genome is represented in blue. K30lambda2.2 is the minor parental and its genome is represented in red. Recombinant fragments that potentially belong to the minor parental are represented in red. **a.** Recombinants obtained in lineage 1. **b.** Recombinants obtained in lineage 2. **c.** Recombinants obtained in lineage 3.

Supplementary Figure 2. **Evolved variants K74_evo1 and K74_evo4 of K74PH129C2 and differences in the increment of titer per time. a.** Representation of the homology of phage variants (K74_evo1 and K74_evo4) and the ancestor (K74PH129C2) using the R package gggenomes (69). The phages are represented as the annotated coding sequences (CDSs). Functions are represented by different colors specified in the legend. The function of deleted fragments with known functions is indicated with *: *1. SAM-dependent methyltransferase. *2. dGTPase inhibitor. **b.** Graphic representation of the difference in the increment of titer per time for each phage. Calculated as follows: $\Delta Log_{10}$ (titer) = $Log_{10}$ (Ti titer) - $Log_{10}$(Tf titer), being Ti = initial time and Tf = final time. The increment of titer was calculated in 6 hours.

Supplementary Figure 3. **Graphic representation of recombinant fragments of evolved *Mydoviruses*.** The figure illustrates the whole genome alignment of all recombinants of each lineage with parental phages. The genome size bar corresponds to positions in the alignment. K54lambda1.1.1 is the major parental and its genome is represented in blue. The minor parental is an unknown sequence. K45PH128C2 is the second *Mydovirus* included in the phage cocktail and its genome is represented in red. Recombinant fragments that potentially belong to minor parentals are also indicated in red. **a.** Recombinants obtained in lineage 1. **b.** Recombinants obtained in lineage 2. **c.** Recombinants obtained in lineage 3.
